# Mechanosensory stimulation via *Nanchung* expressing neurons can induce daytime sleep in *Drosophila*

**DOI:** 10.1101/829861

**Authors:** Shahnaz Rahman Lone, Sheetal Potdar, Archana Venkataraman, Vasu Sheeba, Vijay Kumar Sharma

## Abstract

The neurogenetic bases of sleep, a phenomenon considered crucial for well-being of organisms has recently been under investigation using the model organism *Drosophila melanogaster*. Although sleep is a state where sensory threshold for arousal is enhanced, it is known that certain kinds of repetitive sensory stimuli such as rocking, can in fact promote sleep in humans. Here we report that orbital motion aided mechanosensory stimulation promotes sleep in *Drosophila*, independent of the circadian clock, but controlled by the homeostatic system. Mechanosensory receptor *nanchung* (*Nan*) expressing neurons in the chordotonal organs mediate this sleep induction - flies where these neurons are either silenced or ablated display significantly reduced sleep induction upon mechanosensory stimulation. Transient activation of the *Nan*-expressing neurons also enhances sleep levels confirming the role of these neurons in sleep induction. Thus, we show for the first time that mechanosensory stimulation promotes sleep even in flies *D. melanogaster* and that it is mediated by proprioceptors.

## Introduction

The phenomenon of sleep is exhibited by organisms across the animal kingdom although our understanding regarding functions of sleep still remains inadequate (Saper et al., 2010; Shaw et al., 2013). Animals shut down many vital behaviours such as feeding and reproduction and are more vulnerable to predators during sleep, yet, the sleeping state constitutes a half to one third of their lifespan (Allada and Siegel, 2008; Siegel, 2009; Shaw et al., 2013). It is believed that circadian clocks determine the timing of sleep, while its quality (intensity) and quantity (duration) is determined by the homeostatic system (reviewed in Shaw et al., 2013). Studies on organisms including fruit flies, worms, zebrafish and other model organisms have revealed that many aspects of sleep are genetically determined (Allada and Siegel, 2008; Crocker and Sehgal, 2010; Shaw et al., 2013). However, there is also evidence to suggest that environmental stimuli like light, temperature, social cues etc., can modulate quality and quantity of sleep in flies (Lamaze et al., 2017; Parisky et al., 2016; Lone et al., 2016; Lone et al., 2012; Donlea et al., 2009; Fritzgerald et al., 2006).

A relatively simple nervous system and ease of genetic manipulations has led to the vinegar fly *Drosophila melanogaster* becoming increasingly preferred in the quest to unravel neuronal circuits underlying many behaviours including sleep (Venken et al., 2011; Goodwin et al., 2009; Simpson, 2009). Over last two decades, we have begun to understand the role of circadian circuitry in modulating sleep and to appreciate the contributions of homeostatic processes and their interaction with the circadian circuitry (Parisky et al., 2008; Yadlapalli et al., 2018; Guo et al., 2016; Shang et al., 2013; Shang et al., 2008; Sheeba et al., 2008; Liu et al., 2016; Donlea et al., 2011). Neuropeptides such as Pigment Dispersing factor (PDF), small-neuropeptide factor (s-NPF), diuretic hormone 31 (DH31) and others serve as critical contributors to sleep properties (Kunst et al., 2014; Shang et al., 2013; Sheeba et al., 2008; Shang et al., 2008; Parisky et al., 2008; Lamaze et al., 2018). Thus, we are systematically unraveling the neuronal circuitry that modulates sleep using *D. melanogaster* (Potdar and Sheeba, 2013).

Sleep is also modulated by age, sex, diet and several environmental conditions in flies (Keene et al., 2010; Linford et al., 2012). In humans, walking during daytime has been shown to influence sleep duration and quality of nighttime sleep (Morita et al., 2011). In addition to the conventional prescription of somnolence-inducing drugs, non-pharmacological aids using cognitive behavioural therapy, which includes, relaxation techniques that can reduce hyperarousal in patients, have been used to treat insomnia (Siebern et al., 2012). Additionally, transcranial magnetic stimulation (TMS) (Massimini et al., 2007), transcranial direct current stimulation (tDCS) (Marshall et al., 2006; Reato et al., 2013), open-loop audio-visual stimulation (AVS) (Tang et al., 2016), acoustic stimulation (Bellesi et al., 2014) have been shown to aid sleep in mammals. Although it is common knowledge that babies in cradles, adults in rocking chairs and passengers in moving vehicles, fall asleep readily, yet, the mechanisms underlying motion-induced sleep remain unclear. Human subjects maintained in a linearly accelerated swing made a faster transition to sleep, exhibiting increased number of rapid eye movements as compared to those who stayed stationary (Woodward et al., 1990). A more recent study which examined daytime sleep (afternoon nap) in human subjects demonstrated that gentle rocking movements enabled participants to quickly transition from waking to sleep and enabled them to experience longer duration <u>n</u>on-rapid <u>e</u>ye <u>m</u>ovement - NREM-stage 2 sleep (Bayer et al., 2011). Yet another study on human volunteers (Perrault et al., 2019) showed that sleep latency is reduced by nighttime rocking and there are fewer arousals compared to the night when volunteers were not subjected to rocking. Increase in slow oscillations associated with consolidated sleep and memory formation was also reported by the study (Perrault et al., 2019). Similarly, the stimulation of vestibular and proprioceptive sensory inputs of mice were also seen to elicit a sustained boosting of slow-wave oscillations and increased density of sleep spindles, both of which are indicators of deep sleep (Kompotis et al., 2019; Bayer et al., 2011). The authors hypothesized that rocking mediated sensory inputs may directly or indirectly impact sleep centers in the brain, however, the underlying neuronal mechanisms remain elusive. Here we provide evidence for an ancient origin for such mechanosensory stimulation induced sleep by demonstrating its existence and underlying pathways in fruit flies *D. melanogaster*. We show that this sleep induction is reversible and independent of the circadian clock but regulated by homeostatic mechanisms and mediated by the mechanosensory receptor nanchung (Nan) expressing neurons located in the chordotonal organs.

## Results

### Orbital motion promotes sleep across multiple fly strains

Based on previous studies of mammals including humans where rocking motion was found to induce daytime sleep (Kompotis et al., 2019; Perrault et al., 2019; Bayer et al., 2011) we set out to examine the effect of sustained orbital motion on flies. Here we report for the first time that gentle orbital motion (Figure 1a) induces sleep in *D. melanogaster*. Female *Canton-S* (*CS*) flies showed higher levels of sleep, when provided <u>O</u>rbital <u>M</u>otion either during <u>D</u>aytime (OMD), or <u>c</u>ontinuously throughout day and night (OMC) compared to controls which were not exposed to orbital motion but remained in the same incubator (Figure 1b-d). In males also, OMD caused a significantly higher daytime sleep as compared to controls (*p*<0.0001; data not shown). Daytime sleep of females is generally significantly lower than males in many fly strains (Huber et al., 2004; Andretic and Shaw, 2005), consequently, daytime sleep induction by orbital motion is more conspicuous and consistent. Therefore, for the most part, we report results on the phenomenon of orbital motion-aided enhancement of daytime sleep in females, unless specified otherwise. Orbital motion of - 80-rpm, 120-rpm showed significant increase (80-rpm: *p*<0.05; 120-rpm *p*<0.01), however 600-rpm could not produce significant impact on sleep increase (*p*=0.21; data not shown). Speeds below 80 rpm were not technically feasible. We found that seven different positions on the shaker platform are equally capable of sleep induction (*p*>0.05 data not shown).

**Figure 1.**
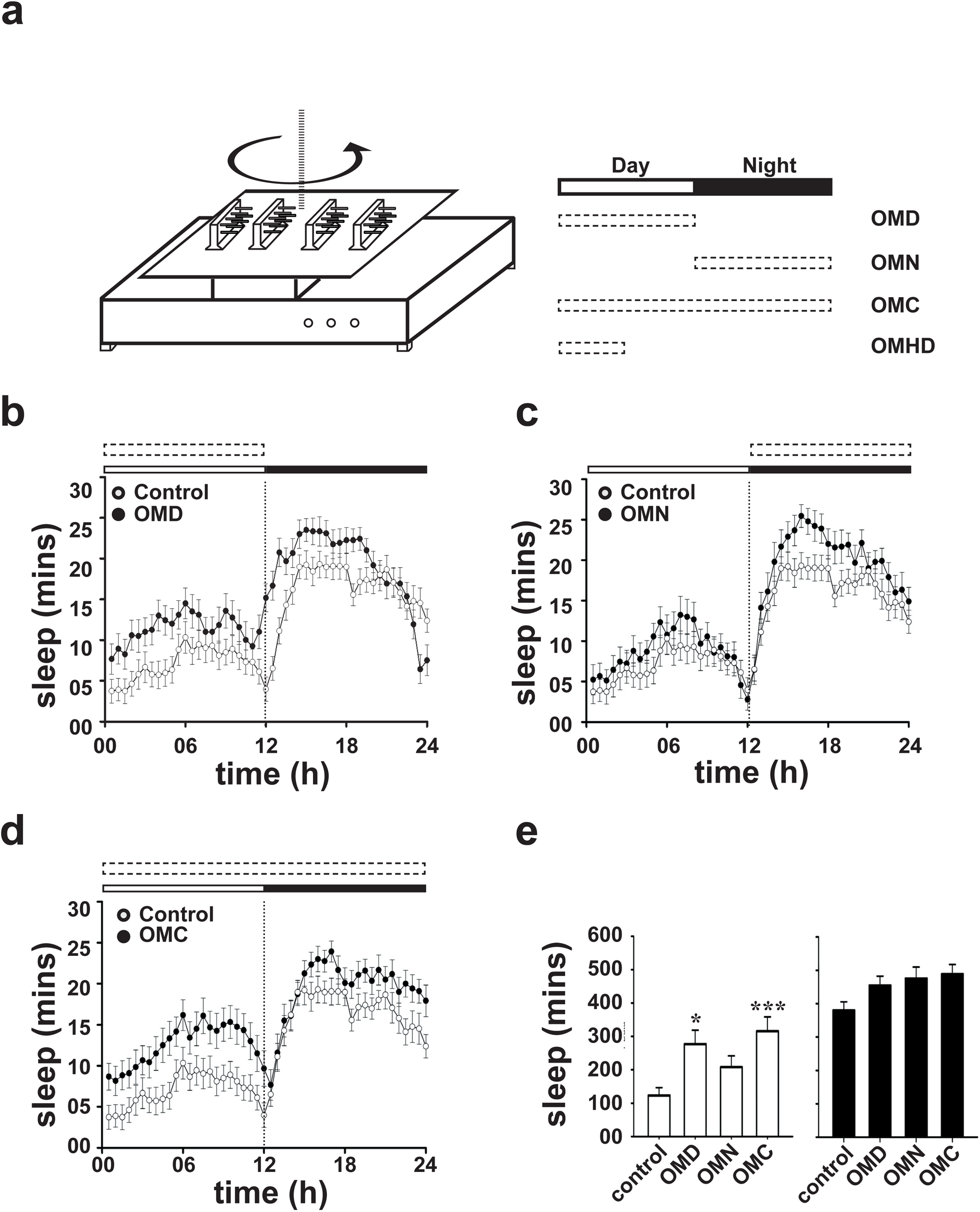
(a-left) Schematic representing experimental protocol used to subject flies to orbital motion. DAM monitors were fixed by double sided adhesive tape on an orbital shaker that rotated the platform in the horizontal plane as shown (120 rpm, unless specified). (a-right) Horizontal white and black bars indicate day and night period under 12:12 h light:dark cycles. Dashed bars indicate the duration for which orbital motion was given – OMD-Orbital Motion during 12h of Daytime; OMN-Orbital Motion during 12h of Nighttime and OMC-Orbital Motion for the entire 24h Cycle. Sleep profile of *CS* flies exposed to (b) OMD, (c) OMN and (d) OMC where (•) indicates treated flies, and (∘) indicate controls. Mean sleep across 3 days averaged across flies (30-min bins) is plotted across time of the day with 12h light phase (white horizontal bar) and 12h dark phase (black horizontal bar). (e) Mean sleep during day (unfilled bars) and night (filled bars) under OMD, OMN and OMC. Error bars are SEM. ANOVA followed by post-hoc multiple comparisons Tukey’s test revealed significant increase (*p* < 0.05) in daytime sleep when exposed to OMD and OMC, whereas OMN has no effect (*p*>0.05). Asterisks denote statistical significance level of *p*<0.05 = *, *p*<0.005 = **, and *p*<0.0005 = ***. Nighttime sleep is not significantly altered in any regime. Sample size varied from n = 24-32 flies.

To uncover any possible strain-specific effect, we examined four different ‘wild-type’ strains of *D. melanogaster- CS, Oregon-R* (*OR*), *w*^*1118*^, *yellow white* (*yw*) and found they also show increase in daytime sleep due to OMD compared to controls (*p*<0.0005; Figure 2a-e). We verified that the orbital motion induced sleep is not due to a general decline in activity since OMD flies showed similar levels of activity per waking min (daytime) as controls (*p*>0.05; Figure 2f). We also observed that orbital motion during daytime induces flies to fall asleep quickly after lights-on, with a statistically significant decrease in daytime sleep latency for most of the strains (*p*<0.0005 for *OR* and *yw*; *p*<0.02 for *CS*; for *w*^*1118*^ *p*=0.16; Figure 2g) while nighttime sleep latency was unaffected (Figure 2h). Thus, we detect a robust phenotype of daytime sleepiness due to OMD that is over and above base line sleep across four fly strains.

**Figure 2.**
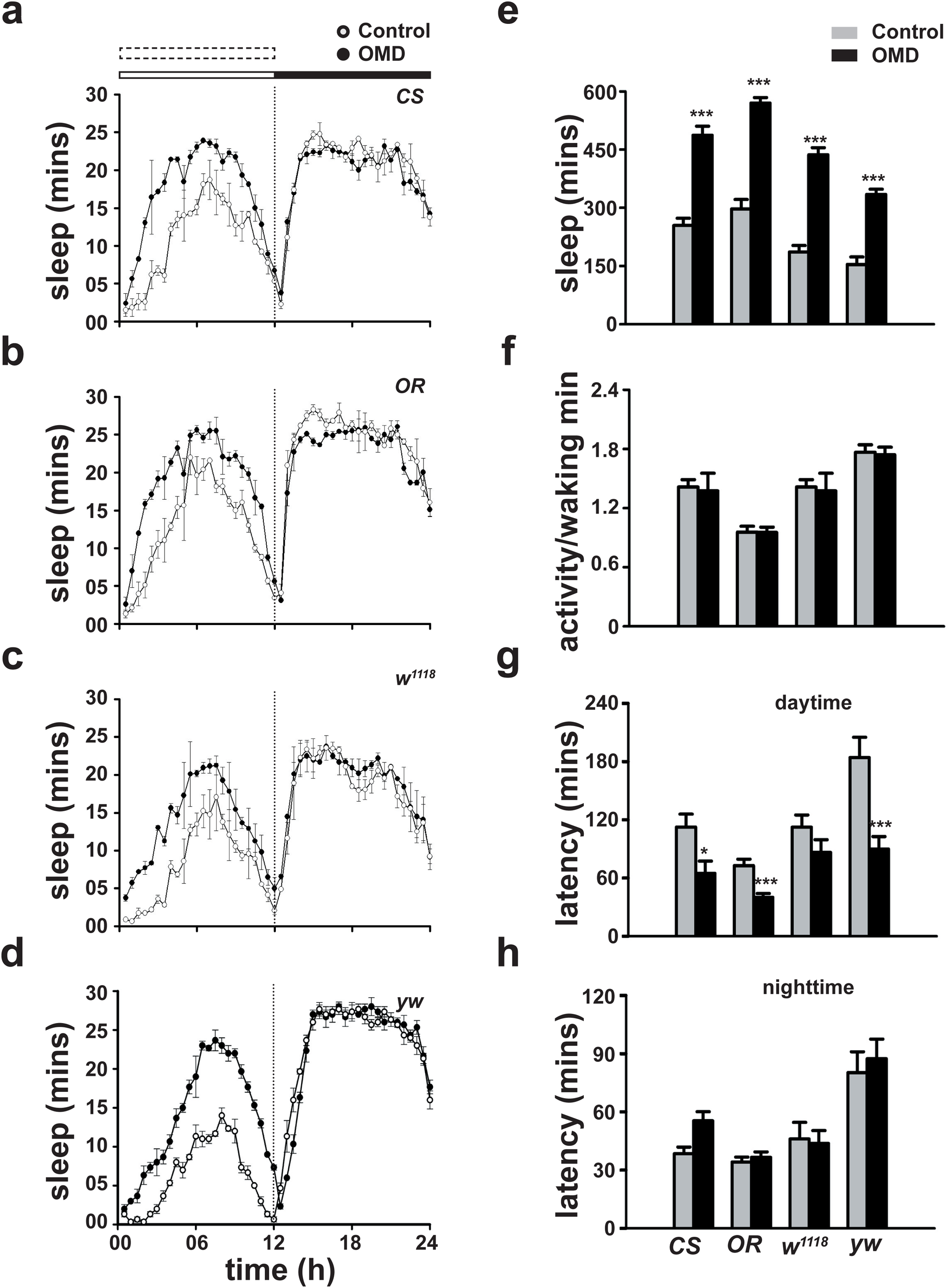
(a-d) Sleep profile of (a) *CS*, (b) *OR*, (c) *w*^*1118*^ and (d) *yw* flies exposed to OMD (•), plotted along with controls (∘). (e) Mean daytime sleep of controls (unfilled bars) and OMD flies (filled bars). ANOVA followed by post-hoc comparisons revealed an increase in sleep (*p*<0.0005) in *CS, OR, w*^*1118*^, and *yw* flies exposed to OMD. (f) Activity per waking min of OMD exposed *CS, OR, w*^*1118*^ and *yw* flies is comparable to controls (*p*>0.05). (g) Daytime sleep latency is smaller for OMD exposed *CS, OR and yw* (*p*<0.0005 for *OR* and *yw, p*<0.02 for *CS*) but remains unchanged in *w*^*1118*^ flies (*p*=0.16). (h) Nighttime sleep latency is not affected (*p*>0.05) in any of the fly strains; All other details same as Figure 1. n = 23-24

### Orbital motion mediated quiescence is “true” sleep with properties of reversibility, homeostatic control and increased arousal threshold

It is possible that the quiescence induced in flies experiencing orbital motion is not true sleep and merely inactivity due to inability to locomote while experiencing such motion. Therefore, we examined whether OMD induced inactivity meets the accepted criteria for sleep in flies, firstly its reversibility, secondly its homeostatic control and thirdly the requirement for a higher arousal threshold (Campbell and Tobler, 1984; Hendricks, 2000; Shaw et al., 2000). To demonstrate reversibility of OMD-induced sleep, both OMD and control groups of flies were subjected to manual physical disturbance by shaking monitors for few seconds during the middle of the day, which is also the peak of daytime sleep in flies, i.e. at Zeitgeber Time 06 (ZT06; by convention ZT00 is lights-ON under a 12:12-h light:dark (LD) cycle) (Figure 3a). Physical disturbance awoke flies in both groups (∼52% and ∼60%) suggesting that this quiescence is reversible. Importantly, flies that experienced OMD fell asleep sooner after physical perturbation as compared to controls (shorter sleep latency *p* < 0.05; Figure 3a,b) suggesting that OMD induced sleep drive has a potent effect on flies even after a recent experience of physical disturbance. It also suggests that orbital motion is effective in promoting sleep at times other than ZT00. We also tested this directly by initiating OMD at ZT04 instead of at ZT00 and similar results were observed for sleep levels and latency (*p*<0.001; Data not shown).

**Figure 3.**
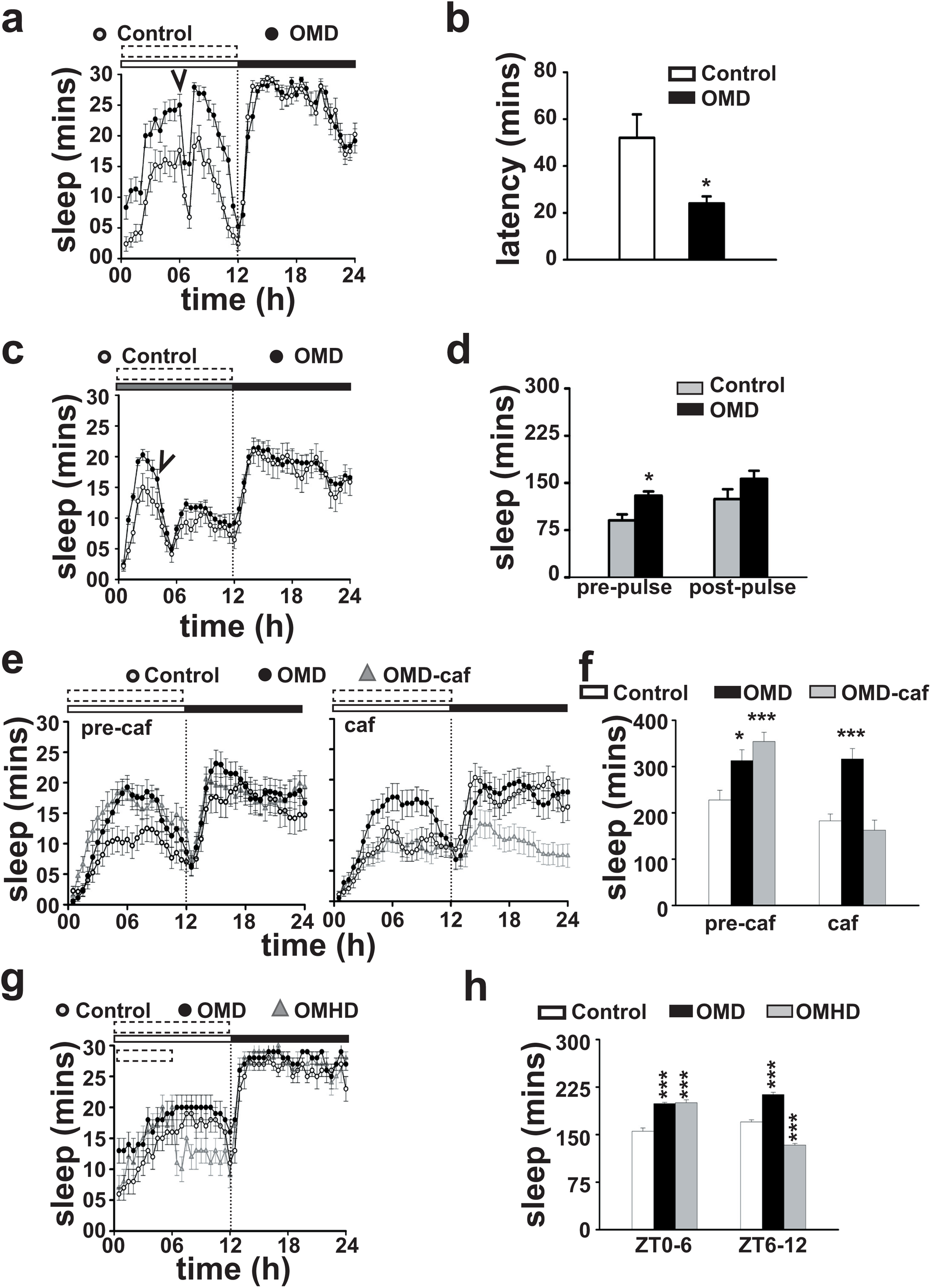
(a) Sleep profiles of *CS* flies with OMD and controls, when physical disturbance was given at ZT06. (b) Sleep latency post physical disturbance at ZT06 of flies from (a) showing OMD flies fall asleep sooner than controls (*p*<0.05). (c) Sleep profile of flies exposed to OMD from CT00 to CT12, and received light pulse of 500-lux for 10-sec on the first day in DD at CT04. (d) Total sleep of flies experiencing OMD from ZT00 to ZT04 is significantly higher (*p*<0.05) than controls. After light pulse, sleep in OMD flies is not different from controls (*p*=0.22). (e-left) Sleep profiles pre-treatment with caffeine (pre-caf) show three sets of *CS* flies where OMD (•) and OMD-caf (▴) show significant increase (*p*< 0.05) in comparison to control (∘). (e-right) While OMD flies (not fed on caffeine) show higher sleep levels than controls, sleep levels of flies fed with caffeine (OMD-caf) fall equal to control levels. (f) Mean daytime sleep of control (white bar) OMD (black bar), OMD-caf (grey bar), showing significant increase (*p* < 0.05) in sleep in response to OMD before caffeine treatment (pre-caf) whereas following caffeine treatment (caf), OMD flies continue to show increase (*p* < 0.0005) in sleep, however OMD-caf flies show similar sleep levels as controls (*p* = 0.98). (g) Sleep profiles of OMD and OMHD (<u>O</u>rbital <u>M</u>otion for <u>H</u>alf-<u>D</u>ay between ZT00-06 only) flies plotted along with controls. Mean sleep of flies in (h) showing significant increase (*p*<0.0005) in sleep in OMD and OMHD flies during the first half of the day (ZT00-ZT06), while in second half of the day (ZT06-ZT12) OMHD flies sleep significantly lesser (*p*<0.0005) than controls whereas OMD flies continue to show significant increase (*p* < 0.0005).

We tested whether orbital motion can induce sleep in absence of light by first subjecting flies to OMD under LD 12:12 for 4 days following which they were placed in constant darkness and flies continue to receive OMD for 12h of subjective day. We found that OMD treated flies exhibited higher sleep levels even during ‘subjective daytime’ (Figure 3c, d, pre-pulse sleep). When exposed to a brief (10-sec) light pulse of 500-lux, 4h after onset of OMD (Circadian Time 04 or CT04) (Figure 3c, arrowhead), similar fraction of OMD flies awoke as controls (Figure 3c,d) (54 % of controls and 60% of OMD flies), which further confirmed that orbital motion-induced quiescence is reversible. Prior to light pulse, OMD flies exhibited significantly higher daytime sleep as compared to controls (*p* < 0.05; Figure 3c,d), however, following light pulse, this difference was not statistically significant (*p*=0.22; Figure 3d). This may be due to overall reduction in sleep for both controls and treated flies brought about by the pulse of light probably due to impact on circadian and sleep parameters.

Like humans, flies are also susceptible to the arousal promoting effects of caffeine (Shaw et al., 2000). We asked whether the OMD is reversible by caffeine. Out of three groups of flies, two received OMD while the third group served as control. Both sets of OMD exposed flies slept significantly more than controls (*p*<0.05; Figure 3e-left panel) on the day prior to caffeine treatment (pre-caf). On the next day, immediately after lights-ON, one of the OMD exposed groups was fed caffeinated food (1-mg/ml, OMD-caf), while the other two groups (OMD and controls) were transferred to fresh food medium without caffeine (Figure 3e-right panel). Flies which received caffeine along with OMD, slept significantly less during daytime as compared to those exposed only to OMD (*p*<0.0005) but not less than the controls (*p*=0.98) (Figure 3f). This suggests that orbital motion-induced sleep can be blocked by pharmacological intervention, emphasizing reversibility of sleep induced by orbital motion and that inactivity is not due to the orbital motion preventing locomotion of flies. Expectedly, nighttime sleep of flies fed with caffeine during the day, was also lower than the other groups possibly due to the persistence of arousal-promoting effect of caffeine into the night.

To examine whether orbital motion-induced sleep can be modulated by homeostatic pathways we divided flies into three groups, the first group was subjected to orbital motion for 12 h during daytime (ZT00-ZT12, OMD), the second group only during the first half of daytime (ZT00-ZT06, Orbital Motion during <u>H</u>alf <u>D</u>ay, OMHD) and the third group served as control. During the first half of the day there was a statistically significant increase in sleep in both groups exposed to orbital motion (OMD and OMHD; *p*<0.0005; Figure 3g,h) as compared to controls. However, during the second half of the day, OMHD group of flies slept significantly lesser than OMD and even lower as compared to controls (*p*<0.0005; Figure 3g, h). This is suggestive of a ‘negative sleep rebound’ during the second half of the day, possibly due to accelerated decline in sleep drive that is believed to occur via homeostatic processes (Hendricks et al., 2000; Shaw et al., 2000). Taken together, we demonstrate that most of the signature features of sleep are conserved in flies exposed to OMD suggesting that OMD-mediated quiescence is in fact “true” sleep.

### Orbital motion induced sleep is independent of circadian clock genes period and Clock-

Circadian clocks are known to influence the timing of sleep (Borbely, 1982; Borbely et al., 2016). Moreover, we saw that the behaviour of OMD induced sleep persists even under constant darkness during subjective day (Figure 3c), so we asked if the circadian clock is required for this sleep-induction. We subjected flies carrying loss-of-function mutation for the *period* (*per*^0^) and *clock* (*Clk*^Jrk^) genes to OMD, and found that flies exposed to orbital motion showed a statistically significant increase in daytime sleep (*p*<0.001 for *per*^0^ and *p*<0.0005 for *clk*^jrk^; Figure 4b) as compared to controls, while their activity counts per waking min did not differ. This suggests that core circadian clock genes *period* and *Clock* are not necessary for OMD-induced sleep.

**Figure 4.**
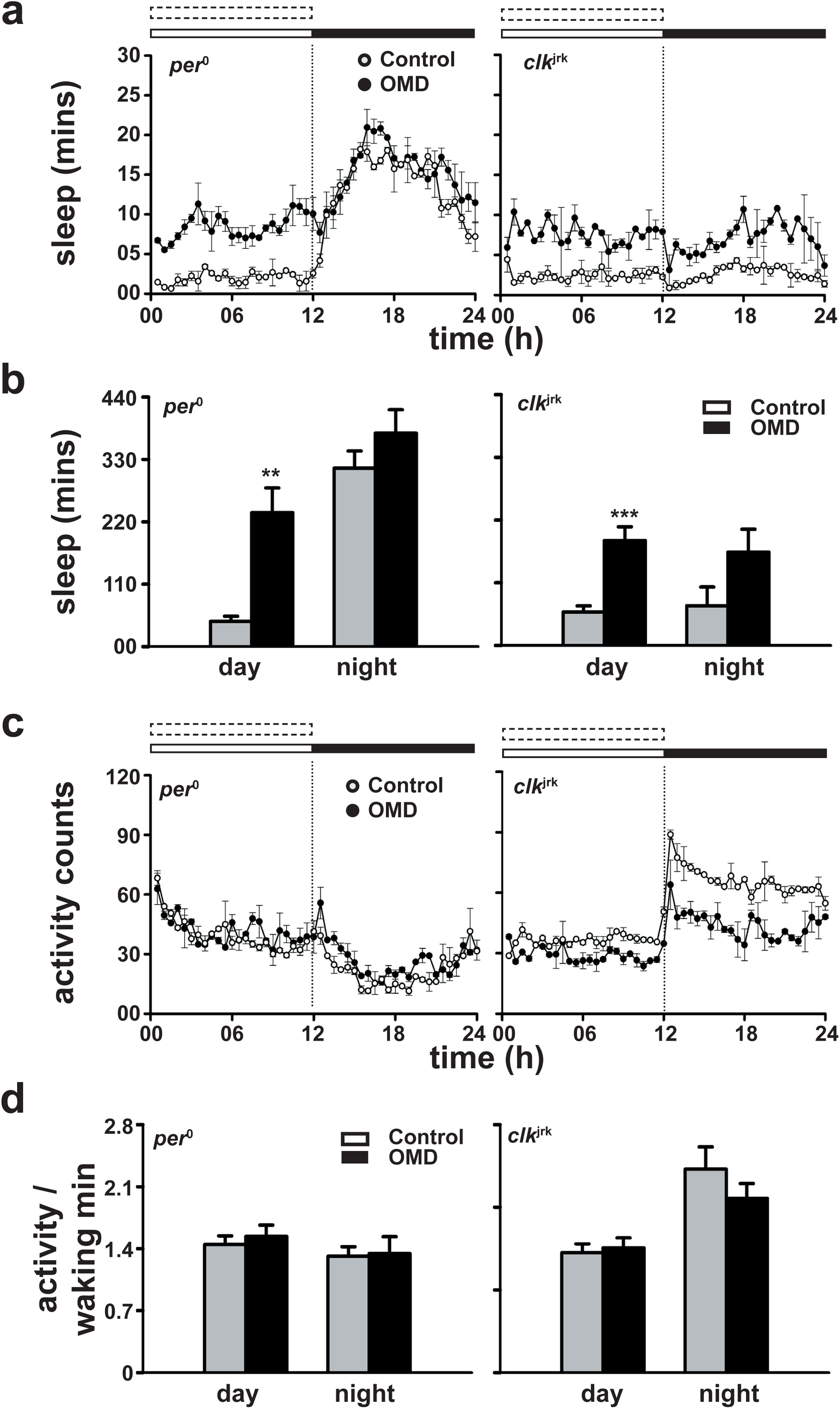
(a-b) Sleep profiles of loss of function mutants of (a) *period* (*per*^0^) and (b) *Clock* (*Clk*^Jrk^). Mean daytime sleep of *per*^0^ and (d) *clk*^jrk^ flies with OMD is higher than controls (*t-test, p*<0.001 for *per*^0^ and *p*<0.0005 for *clk*^jrk^) whereas nighttime sleep is unaffected. (e) Activity profiles of *per*^0^ and (f) *clk*^jrk^. (g) During daytime, activity counts/waking min of *per*^0^ and (h) *clk*^jrk^ flies subjected to OMD did not differ from their controls (*p*>0.05). Other details same as in Figure 1. n ≥13

### Mechanosensory signals via Nanchung expressing neurons mediate orbital motion aided sleep induction independent of other sensory modalities such as vision and olfaction

The mechanosensory system helps organisms to integrate mechanical information to guide their locomotor activity (Tuthill and Wilson, 2016) and locomotion is inhibited during sleep. To examine how orbital motion may transduce signals to the sleep circuit, we examined various mechanosensory signaling pathways. Fruit flies *D. melanogaster* have the ability to sense and respond to touch via specialized mechanosensory centers, the chordotonal organs (Kim et al., 2003), which have been also shown to relay temperature and vibration signals to circadian clocks via yet unknown mechanisms (Sehadova et al., 2009; Simoni et al., 2014). Additionally, mechanosensory neurons present on the legs help flies to avoid aversive conditions by inducing walking behaviour (Ramdya et al., 2014). The chordotonal organs express mechanoreceptor *Nanchung* (*Nan*), which on the basis of sequence similarity is predicted to be an ion channel subunit, similar to vanilloid-receptor-related (TRPV) channels and is activated by forces ranging from osmotic to mechanical pressures (Kung, 2005). These channels also respond to changes in gravity, sound and humidity levels (Kamikouchi, 2009).

We found that flies carrying loss-of-function mutation in the *nan* gene (*nan*^dy5^) (Kim et al., 2003) slept similar (*p*=0.70, Figure 5a-c-) to heterozygote *nan*^*dy5*^*/+* suggesting that reduced *nan* levels does not hamper sleep induction by orbital motion. Since orbital motion is most likely to be transduced to the fly via mechanosensory pathways we targeted the *nan* expressing neurons by expressing the pro-apotoptic gene *hid (nanGAL4>UAShid)*. When we compared sleep change (average sleep of controls with sleep of flies receiving OMD - Δ sleep), *nanGAL4>UAShid* flies showed a significantly reduced response to OMD as compared to their parental controls (*p*<0.0005 for *nanGAL4/+* and *p*<0.01for *UAS/+*; Figure 5d-g). In a separate experiment, we blocked synaptic signaling in *nan*-neurons using an active form of tetanus toxin (*nanGAL4>UAStnt(ac)*) along with controls expressing an inactive form (*nanGAL4>UAStnt(inac)*). Δ sleep in *nanGAL4>UAStnt(inac)* is significantly more (*p* < 0.01, Figure 5h-j) than *nanGAL4>UAStnt(ac)*. These results suggest that orbital motion induces sleep via sensory input received via *Nan*-neurons. We have found similar results when nan expressing neurons were silenced by *UASdORCKC1*. Sleep is increased in *nanGAL4<UASdORKNC* flies subjected to OMD (*p*<0.05), however in *nanGAL4<UASdORKC* flies, there was no significant change in sleep when flies were subjected to OMD (*p* = 0.88; data not shown).

**Figure 5.**
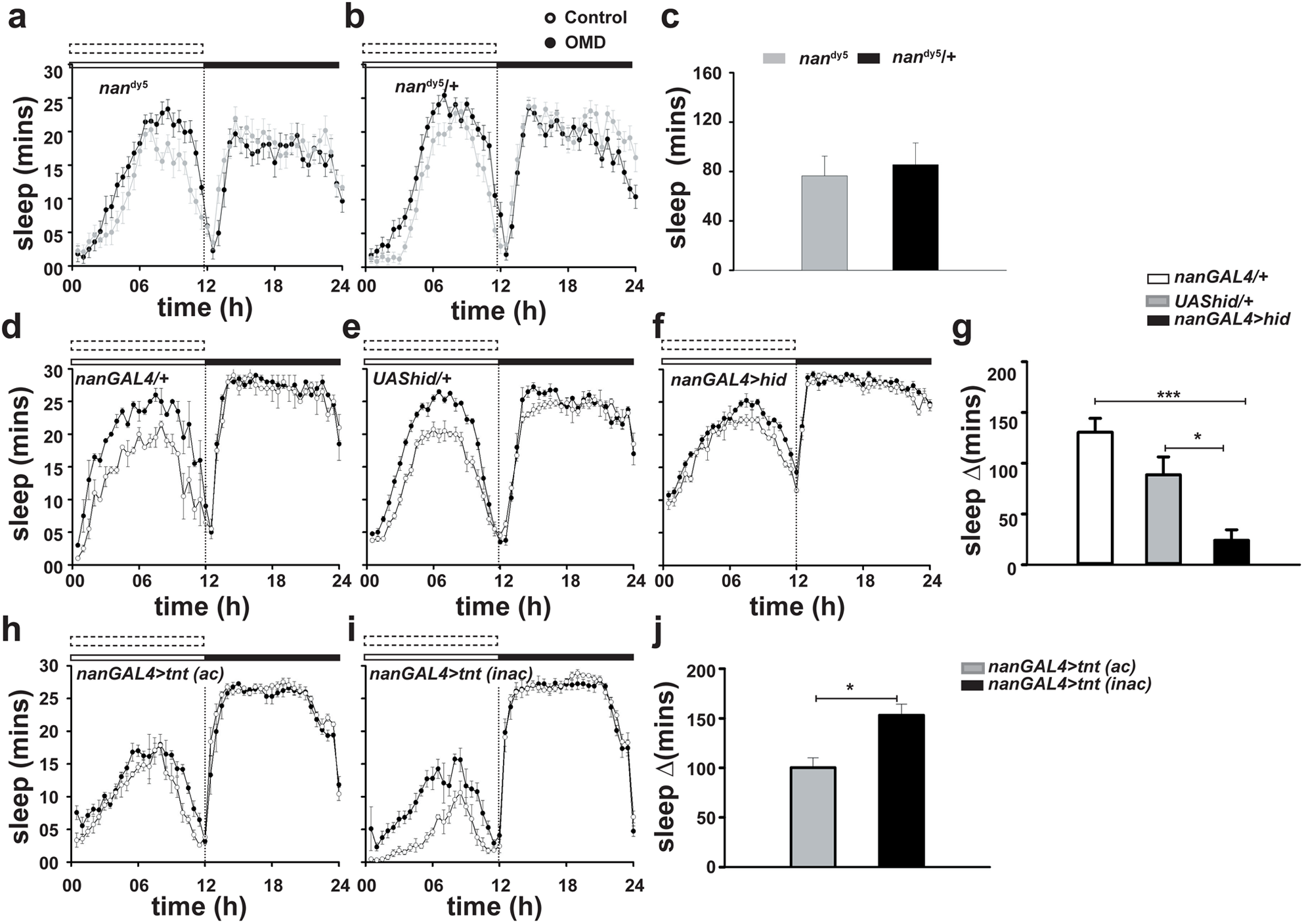
(a-b) Sleep profiles of (a) *nan*^*dy5*^ and (b) *nan*^*dy5*^ /+ flies. (c) Change in daytime sleep levels compared to untreated controls (Δ sleep) of *nan*^*dy5*^ null mutants flies is similar (*p*=0.70-*t*-test) to heterozygote *nan*^*dy5*^/+ controls. (d-f) Sleep profiles of (d) *nanGAL4/+*, (e) *UAS/+* controls, and (f) *nan* ablated *nanGAL4/UAShid* flies. (g) Sleep of nan-ablated neurons (black bar) is significantly reduced (*p* <0.0005 for *GAL4* and *p* < 0.01 for *UAS*) compared to parental controls (*GAL4* (white) and *UAS* (grey)) (h-i) Sleep profiles of (h) flies with electrically silenced nan neurons, *nanGAL4/UAStnt* (ac=active), and (i) control *nanGAL4/UAStnt* flies (inac=inactive). (j) sleep is significantly lower (*p*<0.01) in *nanGAL4/UAStnt* (*ac*) than *nanGAL4/UAStnt* (inac). n ≥16

Since mechanosensory cues are also perceived by auditory and tactile systems we examined mutant flies for the gene *nompc2*, an ion channel essential for mechanosensory transduction, known to have defects in the receptors of tactile bristles (Walker et al., 2000) and found *nompc2* flies exposed to OMD slept significantly more than controls (*p*<0.0001; Figure 6a), which suggests that mechanosensory cues emanating from auditory and tactile organs are not involved in orbital motion mediated sleep-induction, and that mechanosensory signals from parts of the chordotonal organs other than those responsible for auditory signals are likely to be involved in this sleep phenotype. Because mechanosensory cells are also present in the antennae, we examined the role of mechanosensory cues received by the antennae by surgically removing both the antennae of *CS* flies (*CS ant*^*-*^). We found that antennae-less flies also exhibit sleep induction due to OMD (*p*<0.005 for controls and *p*<0.05 for *CS ant*^*-*^ Figure 6b), which suggests that mechanosensory signals from the antennae are not necessary for this sleep phenotype.

**Figure 6.**
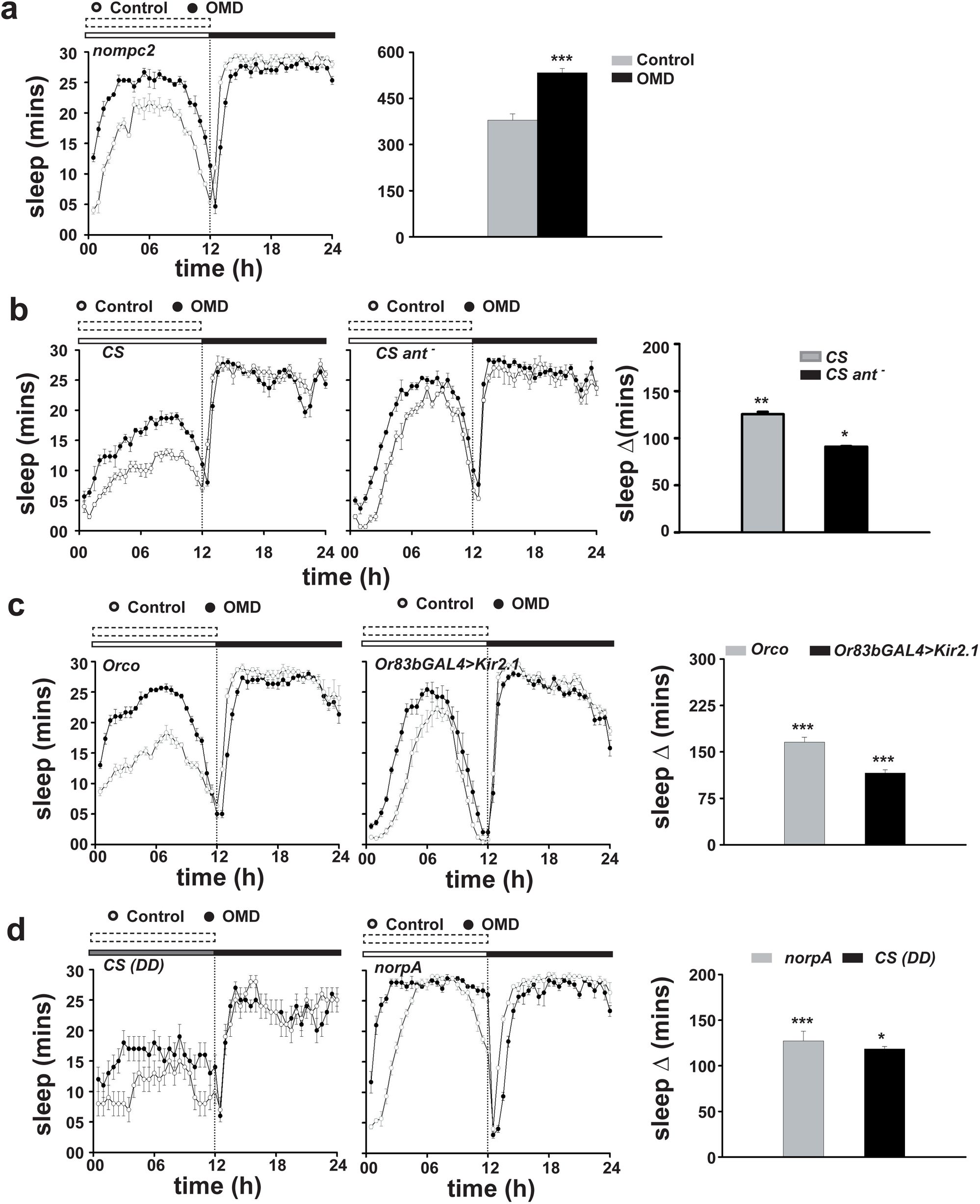
(a-left) Sleep profiles of *nompc2* flies. (a-right) Daytime sleep is significantly increased (*p*<0.0001-t-test) in *nompc2* flies in response to OMD compared to controls. (b-left) Sleep profiles of *CS* flies with antennae intact or (b-middle) surgically removed. (b-right). Sleep of intact flies and flies without antennae is increased in response to the OMD (*p*<0.005 for controls and *p*<0.05 for *CS ant*^*-*^). (c-left) Sleep profiles of *Orco*, and (c-middle) *Or83bGAL4>UASkir2*.1 flies. (c-right) Daytime sleep of *Orco* and *Or83bGAL4>UASkir2*.1 flies show statistically significant increase (*p*<0.0001 –*t* test) in response to OMD compared to controls. (d-left) Sleep profile of *CS* flies in DD (CS-DD) and (d-middle) *norpA* flies subjected to OMD. (d-right) Daytime sleep of CS flies is significantly greater in response to OMD compared to controls (*p*<0.05-t-test). Similarly, *norpA* flies show increase in sleep (*p*<0.0001-*t*-test) when exposed to OMD as compared to controls. n ≥16

Orbital motion, could in principle, have some direct/indirect effect on sensory systems such as olfaction, vision and audition, which in turn may affect sleep circuitry. A previous study on rats has shown that olfactory stimulation can induce slow wave activity (Fontanini and Bower, 2006), while cutaneous stimulation in cats resulted in synchronised brain activity akin to sleep (Pompeiano and Swett, 1962). A more recent study on human subjects showed that acoustic inputs by way of brief auditory tones (0.5 – 4 Hz) induced slow wave activity with features very similar to natural sleep (Tononi et al., 2010). To test for such effects we blocked olfactory and visual inputs to flies while they were subjected to OMD. We tested null mutants of *Orco* (*Or83b*^o^), which are defective in their olfactory ability for most odors (Larsson et al., 2004) and found that mutant flies subjected to OMD show increased (*p*<0.0001; Figure 6c) daytime sleep as compared to controls. Further, when we silenced the *Or83b*-expressing olfactory neurons by expressing potassium channels (*UASkir2*.*1*), flies exposed to OMD showed increased sleep (*p*<0.0001; Figure 6c-e) as compared to controls, which suggests that olfactory signals do not play any role in mechanically stimulated sleep induction.

To examine the role of vision, we subjected *CS* flies to LD cycles for three days and then transferred them to DD along with orbital motion for the first 12 hours under darkness and estimated sleep levels during subjective day. We found a statistically significant increase (*p*<0.05; Figure 6f,h) in daytime sleep in OMD flies compared to controls even under darkness, which suggests that visual cues are not needed for sleep induction. We also found that *norpA* mutants, known to have defective vision (Bloomquist et al., 1988) also showed sleep induction when exposed to OMD (*p*<0.0001; Figure 6g,h). Taken together these results suggest that orbital motion aided sleep induction can occur even under sub-optimal performance of sensory modalities such as olfaction, vision, hearing and tactile sensation.

### Transient activation of Nan-neurons induces sleep in absence of orbital motion

Since our results thus far suggest that *nan*-neurons transduce mechanosensory stimuli produced by orbital motion to sleep centers. We silenced Nan neurons transiently by over expressing *UASshibire*^*ts*^. At lower temperature of 21°C, sleep change (Δ sleep) of *nanGAL4 /UASshibire*^*ts*^ is similar (*p* = 1.0, Figure 7a, c) to controls during daytime and nighttime. When we increased temperature to 28 °C, silenced flies (*nanGAL4 /UASshibire*^*ts*^) do not show sleep induction compared to undisturbed controls as evident by the significantly lower Δ sleep (*p* < 0.0005 for daytime and *p* < 0.05 for nighttime). We asked whether transient activation of *Nan*-neurons alone can induce sleep in the absence of orbital motion. We expressed temperature gated cation channel *dTRPA1* under the *nanGAL*4 driver and found that at sub-threshold temperatures (21°C) *nanGAL4*/*dTRPA1* flies sleep as much as *nanGAL4*/*+* and *dTRPA1/+* controls (*p*> 0.05; Figure 7d, e). However, when the temperature was raised to 28 °C (activation temperature of dTRPA1, Viswanath et al., 2003), and *Nan*-neurons are expected to enhance firing frequency (Hamada et al., 2008; Tang et al., 2013), sleep change is significantly higher in *nanGAL4*/*dTRPA1* flies as compared to *nanGAL4*/*+* (*p* < 0.05) and *dTRPA1/+* (*p* < 0.005) controls (Figure 7d, e), which suggests that electrical activity of *Nan*-neurons contributes to orbital motion induced sleep.

**Figure 7.**
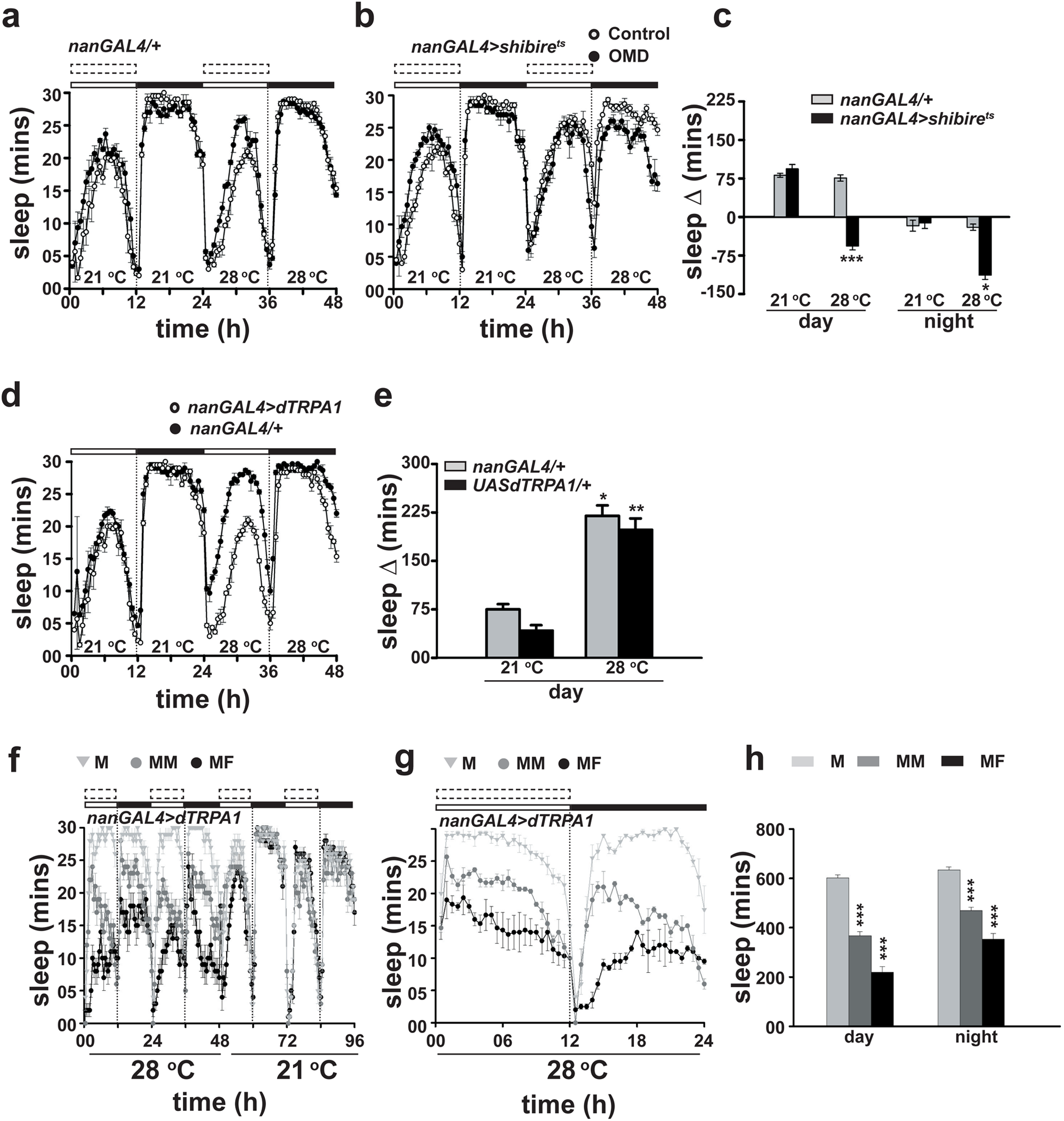
(a-b) Sleep profiles of control (a) *nanGAL4/+* and (b) *nanGAL4/UASshibire*^ts^ flies at 21 °C and 28 °C. (c) Daytime sleep shows similar increase (*p*=0.95, ANOVA followed by Tukey’s test) in response to OMD in both the genotypes at 21 °C. At 28 °C silenced show statistically significant difference (*p*<0.0005) in change in daytime sleep. At 28 °C, nighttime sleep is also reduced (*p*<0.05) in silenced flies compared to controls. (d) Sleep profiles of control (*nanGAL4/+*) and *nan* activated (*nanGAL4/UASdTRPA1*) flies. (e) Daytime sleep in two genotypes does not differ at 21°C, however, when temperature is increased to 28 °C, *nanGAL4/UASdTRPA1* flies show increase in sleep (*p* < 0.05 for *nanGAL4*/*+* and and *p* < 0.005 for *dTRPA1/+* -ANOVA followed by Tukey’s test). (f) Sleep profiles of of *nanGAL4/UASdTRPA1* males cohoused with *CS* females, across two days. Temperature was 28 °C for the first two days and then isolated males (without partners) were returned to 21 °C for two days. (g) average sleep profile at 28 °C (h) Sleep bar graph of *nanGAL4/UASdTRPA1 in* different combination (M, MM, and MF, with F being *CS* instead of *nanGAL4/UASdTRPA1*). Daytime sleep of *nanGAL4/UASdTRPA1* show decrease in sleep compared to controls kept solitary (M) or in group of two (MM). (h) Day as well nighttime sleep in *nanGAL4/UASdTRPA1* male and *CS* female is significantly decreased (*p*<0.0001) compared to those in *nanGAL4/UASdTRPA1* males maintained in pairs or solitarily.. All other details same as in Figure 1. n ≥16

Since it is likely that the activation of *Nan*-neurons using dTRPA1 could cause them to fire at very high rates that could lead to paralysis which we would not be able to distinguish from sleep we carried out another study using a behavioural means to perturb sleep. Male flies become hyperactive in the presence of females and show reduced sleep (Fuji et al., 2007; Lone and Sharma, 2012) we used this behavioural paradigm to disrupt sleep in *nanGAL4*/*dTRPA1* males, where we expect the transient activation of *Nan*-neurons to induce sleep at 28 °C. We paired males of *nanGAL4*/*dTRPA1* with *CS* females and found that at 28 °C the male-female couples show decreased sleep as compared to male-male pairs, or solitary *nanGAL4*/*dTRPA1* males (*p*<0.0001; Figure 7f-h) confirming the finding that activation of *Nan*-neurons does not cause a paralytic effect in either males or females and its effect is more likely to occur via its action on sleep circuits.

We fed *nanGAL4*/*dTRPA1* flies with caffeine which is known to prevent sleep and cause arousal while simultaneously activating the neurons by raising the temperature to 28 °C. Caffeinated *nanGAL4*/*dTRPA1* flies showed significantly lower sleep as compared to flies fed with sucrose (vehicle) specifically under 28 °C when *nanGAL4* driven neurons are excited (*p*<0.0001; data not shown). This shows that *Nan* neuronal activation does not lead to paralysis and that the sleep induced by transient hyperexcitation is reversible.

### Electrical stimulation of arousal neurons can override orbital motion-induced sleep

Previous studies in *Drosophila* suggest a key role for l-LN_v_s in sleep regulation by maintaining the arousal state (Sheeba et al., 2008; Shang et al., 2008; Shang et al., 2011; Shang et al., 2013; Parisky et al., 2008; Donlea, 2009). We asked whether orbital motion can induce sleep when the l-LN_v_ neurons are activated by examining flies with *c929GAL4* driving *NaChBac* (*c929/NaChBac*, Nitabach et al., 2006), Figure 8b,c). We found daytime sleep of *c929/NaChBac* is lower (*p*<0.005 for *c929/dORK*Δ*-NC* and *p*<0.05 for *c929GAL4*) than controls (*c929/dORK*Δ*-NC* and *c929GAL4*). Further, we transiently activated l-LN_v_s using *dTRPA1* in adults and found that at 28 °C, *c929/dTRPA1* flies showed significantly lower sleep (*p*<0.001, *p*<0.05 in comparison to *c929/*+ and *dTRPA1*/+; Figure 8d-f) as compared to parental controls. However, at sub-activation-threshold temperature of 21 °C all three (*c929/dTRPA1, c929/*+ and *dTRPA1*/+) genotypes showed a statistically significant increase in sleep when exposed to OMD, and there was no difference in sleep between the experimental and control flies (*p* > 0.91; Figure 8f). Thus, arousal signals due to electrical excitation of l-LNvs can override sleep signals induced by OMD.

**Figure 8.**
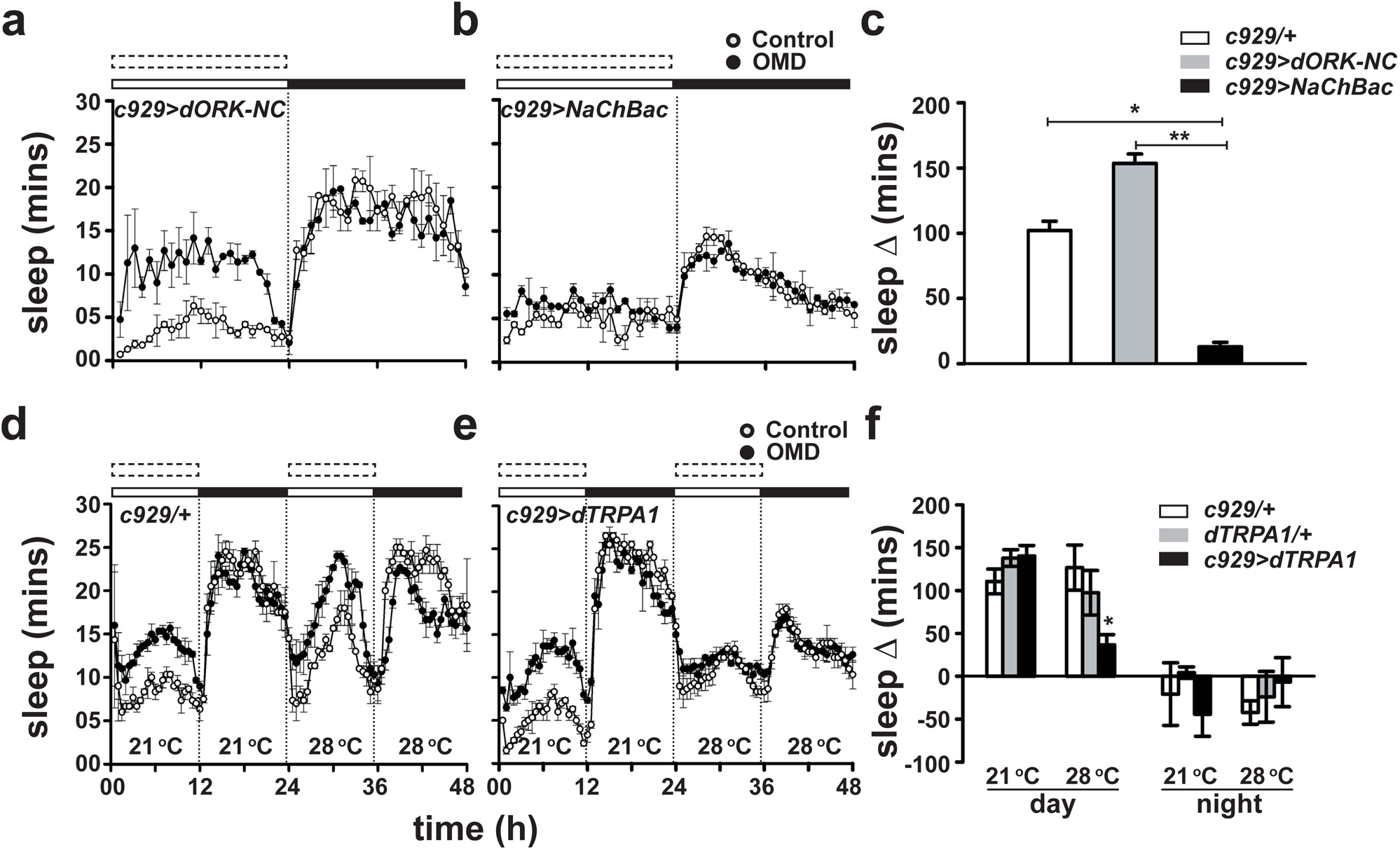
(a-b) Sleep profiles of (a) *c929/dORK*Δ*-NC* and (b) *c929/NaChBac* flies. (c) Daytime sleep of *c929/NaChBac* is lower (*p*<0.005 for *c929/dORK*Δ*-NC* and *p*<0.05 for *c929GAL4*) than *c929/dORK*Δ*-NC* and *c929GAL4*. (d-e) Sleep profiles of (d) *c929GAL4/+* and (e) *c929GAL4/UASdTRPA1* flies. (f) Daytime sleep of all three genotypes viz. *c929GAL4/UASdTRPA1, c929GAL4/+* and *UASdTRPA1/+* show similar (*p*>0.91) at 21°C in response to OMD, whereas at 28 °C, *c929GAL4/UASdTRPA1* flies shows statistically significant decrease in change in sleep (*p*<0.001, *p*<0.05 in comparison to *c929GAL4/*+ and *UASdTRPA1*/+ respectively-ANOVA followed by Tukey’s tests) in comparison to the other two genotypes. All other details same as in Figure 1. n ≥31

## Discussion

Our study examined and found that gentle-movement-induced sleep that humans experience can be replicated in several fly strains and attempted to decipher the neuronal circuits involved. We designed an experimental paradigm involving exposure to orbital motion (Figure 1a) and found that it promotes daytime sleep in fly strains without compromising their overall locomotor activity levels (Figure 2) and that this phenotype is especially evident in females which otherwise exhibit significantly lower daytime sleep levels. Local field potentials in the female fly brain were previously shown to differ between daytime and nighttime sleep (van Alphen et al., 2013) suggesting qualitative differences between them, with daytime sleep deemed to be lighter. Our studies are consistent with previous studies that showed that rocking can induce sleep in humans and mice (Perrault et al., 2019; Kompotis et al., 2019; Bayer et al., 2011). Orbital motion induced quiescence is completely reversible by pharmacological and behavioural interventions (Figure 3) and thus qualifies this criterion for sleep. OMD induced sleep does not require functional circadian clocks (Figure 4) but appears to act via homeostatic processes (Figure 3g, h). Null mutations of core clock genes continue to respond to the OMD and show consistent increase in sleep (Figure 4). We showed that the orbital motion induced sleep relies on mechanosensory *Nan-*neurons present in the chordotonal organs. The role of *Nan* expressing neurons is evident by the fact that mechanically stimulated flies with ablated (*nanGAL4/UAShid*) or silenced (*nanGAL4/UAStnt*) *Nan*-neurons show a significantly reduced sleep induction (Figure 5). The fact that activation of *Nan*-neurons alone causes sleep induction in absence of orbital motion suggests that mechanosensory cues act to activate the chordotonal *Nan*-neurons to promote sleep. Our results clarify that tactile bristles and antennae, considered to be the key components of fly mechanosensory system (Kim et al., 2003; Walker et al., 2000; Effertz et al., 2011; Yan et al., 2013) are not involved in mechanosensory stimulation mediated sleep induction (Figure 6b). Thus, we demonstrate here for the first time that mechanosensory stimulation can lead to sleep induction in fruit flies *D. melanogaster*, and that the underlying mechanisms involve activation of *Nan*-expressing mechanoreceptor neurons located in the chordotonal organs.

Most sensory stimuli are perceived by G-coupled protein receptors, except touch, vibration, and pressure, which are sensed by mechanosensory receptors (Kung, 2005). Mechanosensory neurons respond to such cues by converting them into receptor potentials (Gong et al., 2004) and many organisms have evolved the ability to respond to ligands, which changes the magnitude of such receptor potentials. We believe that differences in the type of motion, and intensity of mechanical stimulation would invoke widely different responses. Rapid to-and-fro motion of about 1000-rpm would cause sleep deprivation (Huber et al., 2004), whereas slow orbital motion of 80-120-rpm would result in sleep induction. A similar feature of no-impact followed by sleep inducing, followed by sleep disrupting effect of sensory stimulation was also hypothesized to explain the effect of increased acoustic stimulation in humans (Bellesi et al., 2014). Stimulation above a certain intensity threshold can effectively induce sleep while stimulation below the threshold is ineffective and stimulation well past the threshold can be disruptive Since increased sensory threshold is a cardinal feature of the sleeping state, the ability of repetitive low amplitude sensory input to induce sleep in an awake animal is counter-intuitive. However, recent studies in mammals including humans (Perrault et al., 2019; Kompotis et al., 2019; Bayer et al., 2011) showed that rocking can enhance the propensity to enter into deep-sleep state. The authors hypothesize that certain types of low-grade sensory stimuli can potentially increase sleep propensity by impinging on brain regions which receive inputs regarding a relaxed or low-stress state or by stimulating sleep centers in the hypothalamus or brain stem region. Alternatively, the authors propose that rocking enhances the degree of synchrony in neural activity in thalamo-cortical regions, which in turn are associated with deep sleep (Bayer et al., 2011).

Non-pharmacological intervention to alleviate sleep disorders are highly desirable since pharmacological agents often have off-target effects and lingering effects persisting into the wakeful state. The fly model for sleep has been particularly useful in unraveling genetic components underlying sleep regulation as well as the neuronal pathways involved, including cellular and molecular components (reviewed in Donelson and Sanyal, 2015). Our finding of a pathway of sensory stimulation that can alleviate sleep levels in this model organism should enable future studies that suggest efficient therapeutic measures to treat sleep defects in a wide range of conditions including circadian sleep phase syndromes, neurodegenerative conditions with associated sleep loss, metabolic syndrome and so on.

## Materials and methods

Flies were collected, sexed, and maintained as virgins in a temperature and humidity-controlled room (25 ± 1°C temperature and 75% relative humidity) under 12:12-h light/dark (LD) cycles, at a density of 30 flies per vial. In most of our experiments, light intensity of 500-lux was used during the light phase of the LD cycles. Four-day old flies were loaded into 65mm tubes with 5% sucrose and cotton at other end by anesthetizing them using carbon-di-oxide. Flies were recorded with the help of DAM (TriKinetics Inc Waltham, USA). These monitors were then placed on an orbital shaker of 13” × 13″ dimension and subjected to orbital motion of 120-rpm (unless specified otherwise). Activity monitors were fixed on the orbital shaker using double-sided adhesive tape, attached at the bottom of the monitor. Activity was recorded in 1-min bin and a minimum 5-min of continuous quiescence was considered as sleep (Hendricks et al., 2000; Shaw et al., 2000), estimated using a sliding time window. In most cases, data collected over a period of 3-days were averaged and analyzed using *t*-test or analyses of variance (ANOVA) followed by post-hoc multiple comparisons using Tukey’s test. To plot sleep and activity profiles, data pooled in 30-min bins and averaged across 3-4-days were used. For most of the experiments flies were given OMD during daytime and were left undisturbed on the shaker during night except for the experiment described in Figure 1, here OMD flies were removed from the shaker at ZT12 and were placed along with controls, whereas OMN flies were placed on the shaker at ZT12. At ZT00, OMD flies were placed on the shaker, while OMN flies were removed and placed along with controls. OMC flies remained on the shaker for the duration of 24 hrs.

For perturbation experiments, at ZT04 or ZT06, experimental flies and control flies were subjected to perturbation by physically shaking the monitors. For light pulse exposure under DD, light of 500-lux was switched on for 10-sec without causing any disturbance to flies. To analyze the effect of caffeine on sleep we used two doses of caffeine (1-mg/ml and 4-mg/ml). The 1-mg/ml dose did not cause fly deaths and therefore we continued with this dose for the rest of the experiments. Sleep change (Δ sleep) was estimated by subtracting mean daytime or nighttime sleep of controls from daytime or nighttime sleep of each OMD fly. For figure 7d,e sleep change was calculated as change in sleep of *nanGAL4>dTRPA1* with respect each parental control. For socio-sexual interaction experiment, activity of flies was recorded in 7-mm tubes at 28 °C instead of 5-mm tubes used for all other experiments. For the temporal silencing and activation experiments by *UASshibire*^ts^ and *UASdTRPA1*, a temperature of 28 °C was used, whereas 21 °C was used as the baseline temperature.

